# The Polo kinase Cdc5 is regulated at multiple levels in the adaptation response to telomere dysfunction

**DOI:** 10.1101/2021.10.20.465143

**Authors:** Héloïse Coutelier, Oana Ilioaia, Jeanne Le Peillet, Marion Hamon, Damien D’Amours, Maria Teresa Teixeira, Zhou Xu

## Abstract

Telomere dysfunction activates the DNA damage checkpoint to induce a cell cycle arrest. After an extended period of time, however, cells can bypass the arrest and undergo cell division despite the persistence of the initial damage, a process called adaptation to DNA damage. The Polo kinase Cdc5 in *Saccharomyces cerevisiae* is essential for adaptation and for many other cell-cycle processes. How the regulation of Cdc5 in response to telomere dysfunction relates to adaptation is not clear. Here, we report that Cdc5 protein level decreases after telomere dysfunction in a Mec1-, Rad53- and Ndd1-dependent manner. This regulation of Cdc5 is important to maintain long-term cell cycle arrest but not for the initial checkpoint arrest. We find that both Cdc5 and the adaptation-deficient mutant protein Cdc5-ad are heavily phosphorylated and several phosphorylation sites modulate adaptation efficiency. The PP2A phosphatases are involved in Cdc5-ad phosphorylation status and contribute to adaptation mechanisms. We finally propose that Cdc5 orchestrates multiple cell cycle pathways to promote adaptation.

## Introduction

In response to DNA damage, cells ensure genome stability by repairing the initial injury. The DNA damage checkpoint (DDC) detects and processes the damage, and arrests the cell cycle to provide time for repair. The coordination between the cell cycle and the repair pathways ensured by the DDC is essential to prevent chromosome segregation with unrepaired damage. When the cell fails to repair the damage, a process called adaptation to DNA damage allows the bypass of the checkpoint arrest and completion of mitosis despite the presence of unrepaired DNA damage, thus inducing genome instability (Galgoczy & Toczyski, 2001, Lee, Moore et al., 1998, Sandell & Zakian, 1993, Toczyski, Galgoczy et al., 1997). In unicellular eukaryotes, adaptation may be viewed as a survival strategy for cells experiencing unrepairable damage that can be asymmetrically segregated into daughter cells or be dealt with in the next cell cycle (Coutelier, Xu et al., 2018, Galgoczy & Toczyski, 2001, Kaye, Melo et al., 2004, Roux, Salort et al., 2021). In mammals though, adaptation may contribute to tumor emergence and progression by evading checkpoint control and promoting genome instability (Gutteridge, Ndiaye et al., 2016, Strebhardt & Ullrich, 2006). While the mechanisms and factors involved in the DDC and repair pathways have been studied for decades, our understanding of adaptation to DNA damage is more limited.

In the model organism *Saccharomyces cerevisiae*, a double-strand break (DSB) is signaled by the phosphatidylinositol-3’-kinase-like kinases (PIKK) Tel1 and Mec1. Mec1 phosphorylates the adaptor protein Rad9 and the kinases Rad53 and Chk1, which are recruited to Rad9. The full activation of Rad53 requires an additional autohyperphosphorylation step. Both Rad53 and Chk1 activation promote the stability of securin Pds1 to prevent chromosome segregation (Agarwal, Tang et al., 2003, Cohen-Fix & Koshland, 1997, Sanchez, Bachant et al., 1999). Phosphorylated Rad53 inhibits the mitotic exit network (MEN) by targeting the Polo kinase Cdc5 (Cheng, Hunke et al., 1998, Sanchez et al., 1999) and triggers a transcriptional response, which includes the direct inhibition of the transcriptional activator Ndd1, controlling the expression of the *CLB2* cluster of mitotic genes (Edenberg, Vashisht et al., 2014, Gasch, Huang et al., 2001, Jaehnig, Kuo et al., 2013, Spellman, Sherlock et al., 1998, Yelamanchi, Veis et al., 2014). When no repair is possible, cells stay arrested for 4 to 16 hrs but then undergo adaptation to DNA damage and alleviate the DDC arrest by returning Rad53 and Chk1 to their unphosphorylated state (Lee, Pellicioli et al., 2000, Pellicioli, Lee et al., 2001), while maintaining the upstream part of the DDC largely intact (Donnianni, Ferrari et al., 2010, Melo, Cohen et al., 2001, Vidanes, Sweeney et al., 2010). Besides DSBs, telomere dysfunction is also perceived by the cell as a persistent DNA damage, and has been widely used as a model to study the DDC and adaptation. Telomere dysfunction can be induced by the conditional loss of Cdc13, an essential factor for telomere protection, which is achieved by placing the *cdc13-1* mutant at restrictive temperature to generate resected telomeres and activate the DDC (Garvik, Carson et al., 1995, Toczyski et al., 1997).

Cdc5 is an essential kinase of the cell cycle regulating a variety of processes in mitosis and cytokinesis (Botchkarev & Haber, 2018). Cdc5’s expression starts at S phase and increases in the later phases of the cell cycle until cytokinesis (Charles, Jaspersen et al., 1998, Cheng et al., 1998, Shirayama, Zachariae et al., 1998), consistent with *CDC5* belonging to the cell-cycle regulated *CLB2* cluster of genes, whose coordinated expression strongly depends on the transcription factor complex consisting of Mcm1, Fkh2 and Ndd1 (coactivator) or Isw2 (corepressor) (Koranda, Schleiffer et al., 2000, Sherriff, Kent et al., 2007, Spellman et al., 1998, Zhu, Spellman et al., 2000). Phosphorylation by the cyclin-dependent kinase 1 (Cdk1) is required to stabilize Cdc5 (Crasta, Lim et al., 2008, Simpson-Lavy & Brandeis, 2011). After mitosis, Cdc5 is ubiquitinated by the anaphase promoting complex (APC) associated with Cdh1 and targeted for proteasomal degradation (Charles et al., 1998). The kinase activity of Cdc5 depends on phosphorylation by Cdk1 and potentially other kinases at several sites, including amino acids T70, T238 and T242 (Mortensen, Haas et al., 2005, Rawal, Riccardo et al., 2016, Rodriguez-Rodriguez, Moyano et al., 2016). Among the several mutants of adaptation that have been identified over the past two and a half decades (Harrison & Haber, 2006, Serrano & D’Amours, 2014), a point mutant allele of *CDC5*, called *cdc5-ad* (for “adaptation-defective”; L251W), is specific for adaptation and does not seem to affect most other functions of Cdc5 in a normal cell cycle (Charles et al., 1998, Rawal et al., 2016, Toczyski et al., 1997). Besides, overexpression of Cdc5 accelerates adaptation, suggesting that Cdc5 plays a promoting role in adaptation (Donnianni et al., 2010, Hu, Wang et al., 2001, Sanchez et al., 1999, Vidanes et al., 2010). However, despite its central role in adaptation, how Cdc5 is regulated in response to DNA damage is not well characterized. In studies on global gene expression in response to alkylating agent methyl methionine sulfonate (MMS) treatment and gamma irradiation, CDC5 transcripts were found to be decreased in a *MEC1*- and *RAD53*-dependent manner (Edenberg et al., 2014, Gasch et al., 2001, Jaehnig et al., 2013). In contrast, Cdc5’s protein level and kinase activity in response to MMS or telomere dysfunction have not been conclusively established as reports differed in their experimental conditions and conclusions (Charles et al., 1998, Cheng et al., 1998, Hu et al., 2001, Maas, Miller et al., 2006, Ratsima, Serrano et al., 2016, Rawal et al., 2016, Vidanes et al., 2010, Zhang, Nirantar et al., 2009).

Here, we investigate the regulation of Cdc5 and the adaptation-defective mutant Cdc5-ad in response to telomere dysfunction. We find that the DDC induces a Mec1/Rad53- and Ndd1-dependent downregulation of Cdc5 expression, which is critical to prevent cells from adapting prematurely. We also identify phosphorylation sites of Cdc5/Cdc5-ad that are involved in the adaptation response to telomere dysfunction. We finally propose that Cdc5 acts in multiple pathways to control adaptation, which might explain the complexity of its regulation in response to telomere dysfunction.

## Results

### Cdc5 protein level decreases in response to telomere dysfunction

To understand how Cdc5 is regulated in response to DNA damage, we induced telomere deprotection by incubating *cdc13-1* cells at the restrictive temperature of 32°C and analyzed C-terminally 3xHA-tagged Cdc5 proteins by western blot at different time points (Fig. 1A). We observed that the amount of Cdc5 decreased from 3 hrs onward compared to time point 0. This result was confirmed in a strain with an untagged Cdc5, using an anti-Cdc5 antibody for western blot (Supplementary Fig. 1A). Cdc5 was degraded by the proteasome as evidenced by its stabilization in the presence of the proteasome inhibitor MG132 in the culture media (Supplementary Fig. S1B). The previously identified APC/C-Cdh1-dependent ubiquitinylation motifs, KEN box and destruction box 1 (Arnold, Hockner et al., 2015, Charles et al., 1998), did not appear to be involved in this degradation (Supplementary Fig. S1C and D).

**Figure 1.**
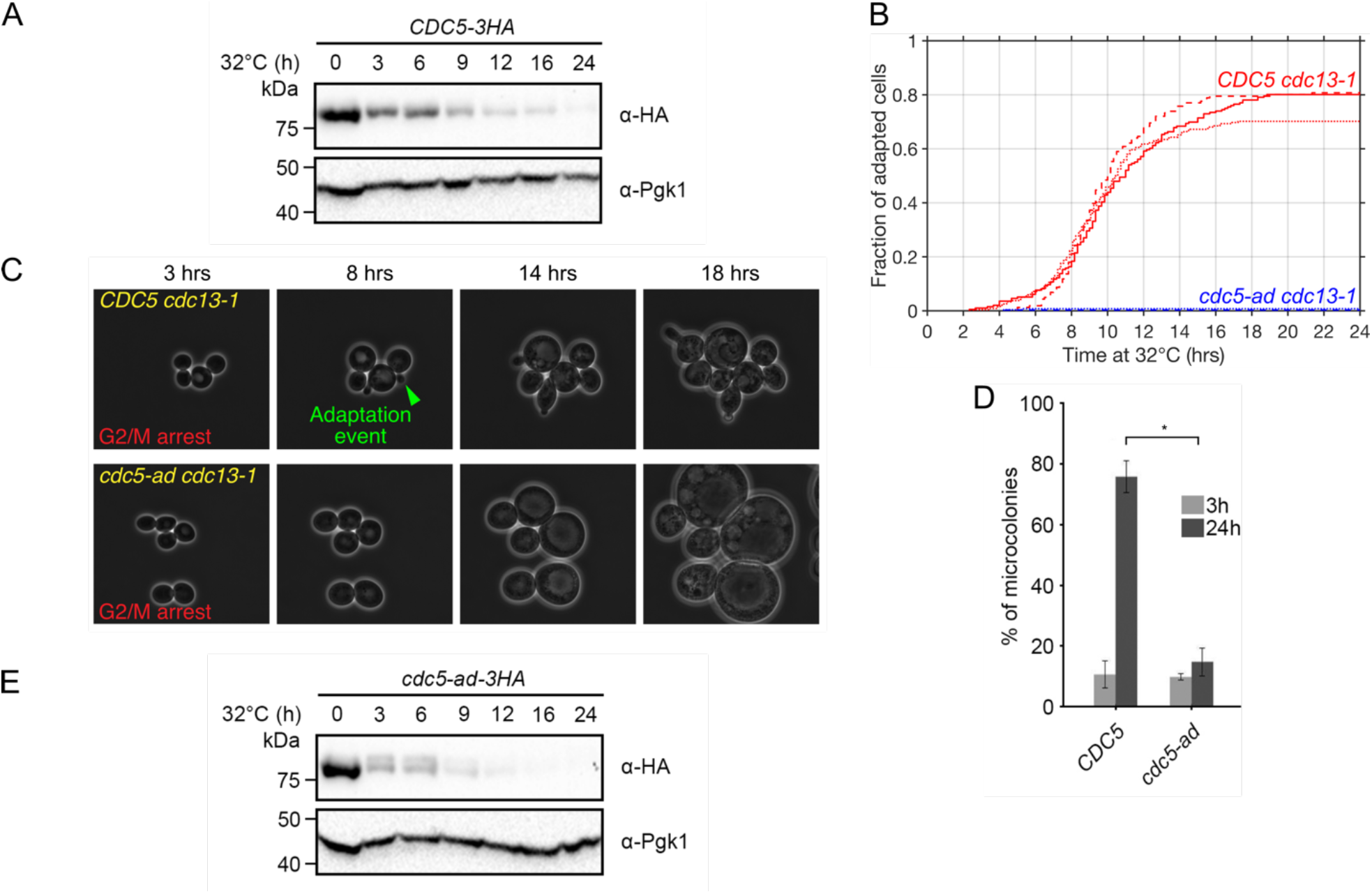
Adaptation response to telomere dysfunction at the cellular and protein level. (**A**) Representative western blot of Cdc5 in a time course experiment after induction of telomere dysfunction by incubating *cdc13-1* cells at 32°C. Pgk1 is shown as a loading control. (**B**) Cumulative curves of the fraction of adapted cells as a function of time after induction of telomere dysfunction, obtained from microfluidic analysis of adaptation events at the single cell level, in *CDC5* and *cdc5-ad* cells. Three independent experiments for each strain are shown in continuous line, dashed line and dots. (**C**) Sequential microscopy images of *CDC5* and *cdc5-ad* cells monitored in microfluidics chambers, used for the analysis shown in (B). (**D**) Microcolony assay measuring the fraction of microcolonies formed by *CDC5* and *cdc5-ad* cells at 3 and 24 hrs. Data are presented as means ± SD of N = 3 independent experiments. n ≥ 150 cells for each condition. **(E)** Representative western blot of Cdc5-ad in a time course experiment after induction of telomere dysfunction by incubating *cdc13-1* cells at 32°C. Pgk1 is shown as a loading control.

We compared the kinetics of Cdc5 degradation with the kinetics of adaptation in the same strain. To achieve a time-resolved detection of adaptation events, we developed a microfluidics-based assay to monitor cells over 24 hrs in time-lapse microscopy with the acquisition of one image every 10 minutes (Fig. 1B and C). Exponentially growing *cdc13-1* cells were loaded into a microfluidic plate as a low-density monolayer in microchambers, set in an incubator at 32°C under the microscope. The chambers were fed with a constant flow of rich media (20 μL/hr for 0.036 μL chambers). Multiple fields of view representing > 150 cells for each condition or strain were monitored in an automated manner. The cells arrested in G2/M within 2-3 hrs (Fig. 1C). Similar to the standard microcolony assay used to assess adaptation (Toczyski, 2006), we used cell rebudding as an indication that an adaptation event occurred. Only the adaptation of an initially arrested cell was counted as one event and we did not count the subsequent cell divisions of adapted cells. In the *CDC5 cdc13-1* strain, the cumulative fraction of adaptation events followed a sigmoid shape with a plateau at 77.0 ± 5.9% (mean ± SD) reached at 16-18 hrs and half of the adaptation events observed at *t*_*50*_ = 9.36 ± 0.30 hrs (mean ± SD) (Fig. 1B). This measurement was consistent with the adaptation percentage obtained by microcolony assay (mean ± SD = 75.7 ± 10.2%) (Fig. 1D), where arrested cells were plated at low density and microcolonies containing more than 3 cell bodies were counted after 24 hrs at 32°C (Toczyski, 2006).

The amount of Cdc5 protein in response to telomere dysfunction was inversely correlated to the kinetics of adaptation events, which was surprising since Cdc5 activity is essential for adaptation. We wondered whether this observation could be explained by the increasing fraction of adapted cells entering the next G1 and degrading Cdc5. To test this hypothesis, we performed the same experiment in the adaptation-deficient *cdc5-ad* mutant, which stays blocked in G2/M. Using our microfluidics assay, we found that virtually no *cdc5-ad* cell underwent adaptation (mean ± SD = 0.6 ± 0.6%) (Fig. 1B). However, the amount of Cdc5-ad proteins decreased in response to telomere dysfunction with a kinetics similar to Cdc5, suggesting that the decrease of Cdc5 protein level was not due to cells entering the next G1, and potentially occurred in G2/M (Fig. 1E). Consistently, we found that in *cdc13-1* cells arrested at 32°C in the presence of nocodazole, which prevents cycling past G2/M, Cdc5 also decreased in quantity at 3 and 6 hours (see below).

### Rad53 inhibits *CDC5* transcription through Ndd1 phosphorylation

*CDC5* belongs to the *CLB2* cluster of genes and their expression depends on the transcription factor Fkh2/Mcm1 associated with the coactivator Ndd1 (Spellman et al., 1998). After DDC activation, Ndd1 is phosphorylated by Rad53 and no longer binds Fkh2/Mcm1, thereby limiting the expression of some of these genes (Edenberg et al., 2014, Yelamanchi et al., 2014). While not all genes in the *CLB2* cluster are repressed through this mechanism in response to DNA damage, the amount of CDC5 mRNAs was found to be decreased (Edenberg et al., 2014, Gasch et al., 2001, Jaehnig et al., 2013). We therefore asked whether the decrease of Cdc5 at the protein level could reflect a repression of *CDC5* transcription through Rad53-dependent phosphorylation of Ndd1.

We thus introduced in our strains a second copy of wild-type *NDD1* or mutant *ndd1-CD-10A* allele (called “*ndd1-10A*” hereafter) expressed from *NDD1*’s native promoter, in which 10 Rad53-dependent phosphorylation sites are mutated (Yelamanchi et al., 2014). We chose to keep the endogenous *NDD1* gene to cover for potential pleiotropic phenotypes of the *ndd1-10A* mutant in cell growth for instance. Since Ndd1 is expected to be excluded from promoters in response to telomere dysfunction, keeping the endogenous *NDD1* gene should not prevent the assessment of the dominant effects of Ndd1-10A recruitment. We then monitored Cdc5 or Cdc5-ad protein levels in these strains after inducing telomere deprotection (Fig. 2A). We found that in contrast to control strains, Cdc5 and Cdc5-ad proteins were fully stabilized in *ndd1-10A* strains, suggesting that the Rad53-dependent repression of *CDC5* and *cdc5-ad* transcription directly translates into a decrease in protein level. Maintenance of Cdc5 and Cdc5-ad protein levels was not due to a loss of checkpoint activation, since Rad53 was hyperphosphorylated in the *ndd1-10A* mutant at restrictive temperature (Fig. 2A).

**Figure 2.**
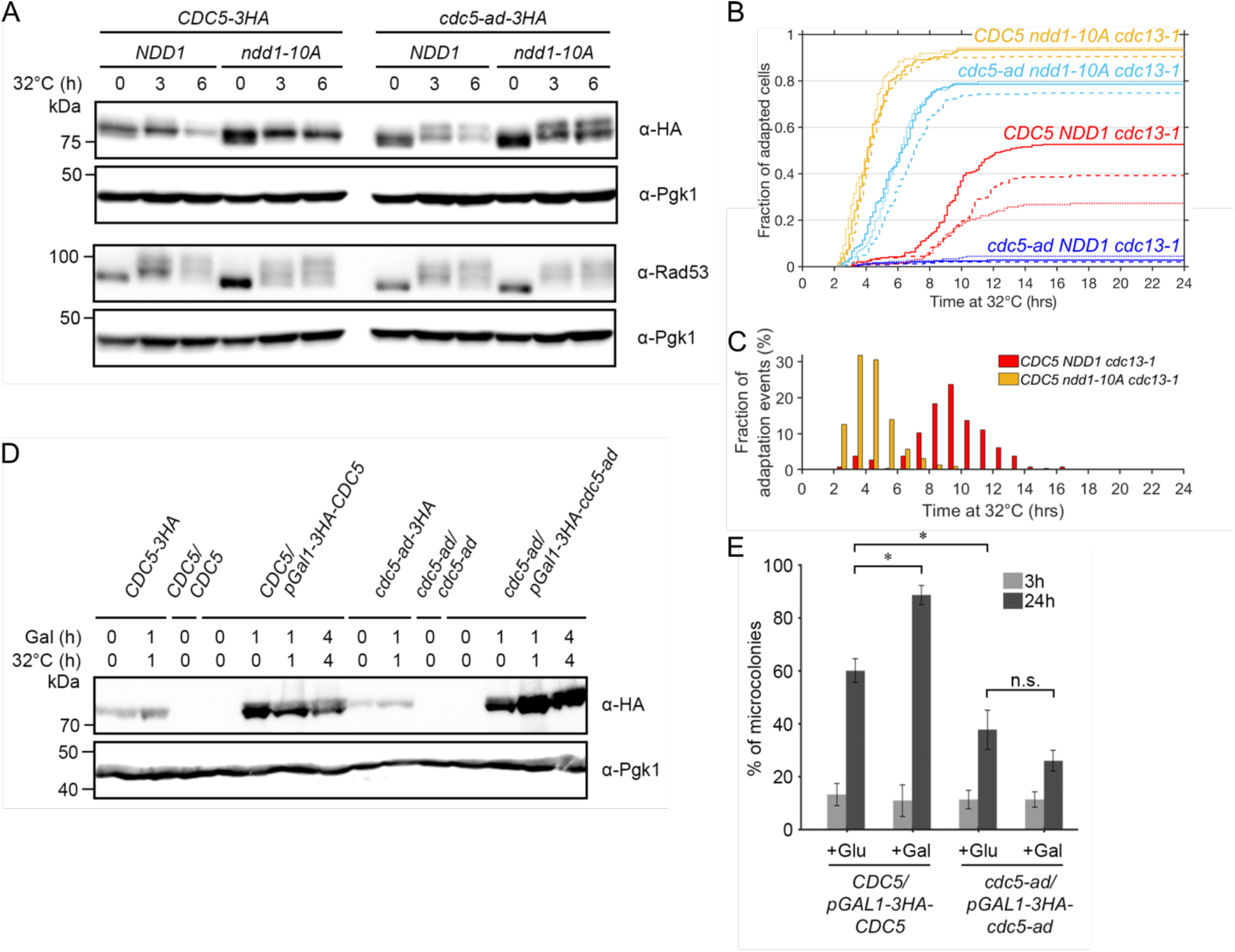
Ndd1 regulates Cdc5 protein level and adaptation kinetics but overexpression of Cdc5-ad does not rescue adaptation defect. (**A**) Representative western blot of Cdc5 and Rad53 in a time course experiment after induction of telomere dysfunction by incubating *cdc13-1* cells at 32°C, in the indicated strains. Pgk1 is shown as a loading control. The presence of the endogenous *NDD1* gene is omitted in the genotype notation for simplicity, for panels (A-C). (**B**) Cumulative curves of the fraction of adapted cells as a function of time after induction of telomere dysfunction, in the indicated strains. Three independent experiments for each strain are shown in continuous line, dashed line and dots. **(C)** Distribution of the timing of the adaptation events for the indicated strains, using data shown in (B). (**D**) Representative western blot of Cdc5 with or without overexpression in the indicated *cdc13-1* diploid and haploid strains. Pgk1 is shown as a loading control. (**E**) Microcolony assay measuring the fraction of microcolonies formed in the indicated strains at 3 and 24 hrs, with or without galactose-induced overexpression of Cdc5/Cdc5-ad. Data are presented as means ± SD of N = 3 independent experiments. n ≥ 150 cells for each condition.

This result further supports the idea that the variation in Cdc5 quantity is a direct response to DDC activation and precedes adaptation.

### The Rad53-Ndd1 signalling pathway regulates the kinetics of adaptation

The role of Cdc5 in adaptation is dose-dependent and overexpression of Cdc5 accelerates adaptation (Donnianni et al., 2010, Dotiwala, Haase et al., 2007, Hu et al., 2001, Vidanes et al., 2010), but this property was previously tested with constitutive expression of *CDC5* at different levels (*e*.*g*. gene copy number or galactose-inducible promoter). We therefore wondered whether adaptation would also be accelerated in the *ndd1-10A* mutant where Cdc5 was not overexpressed but just maintained at its initial level. Adding a second copy of wild-type *NDD1* in the *CDC5 cdc13-1* strain affected the final percentage of adapted cells (mean ± SD = 39.8 ± 12.9%), but not the kinetics (*t*_*50*_ = 9.43 ± 0.57 hrs, mean ± SD) (Fig. 2B). When *CDC5 cdc13-1* cells expressed *ndd1-10A* as a dominant second copy, adaptation was dramatically accelerated (*t*_*50*_ = 4.10 ± 0.06 hrs, mean ± SD) and the total fraction of adapted cells also increased (mean ± SD = 92.3 ± 2.0%) (Fig. 2B). Importantly, we observed that *ndd1-10A* cells arrested in G2/M within the first 2 hrs, indicating that the checkpoint arrest was functional, consistent with Rad53 being hyperphosphorylated upon induction of telomere deprotection in this mutant. Interestingly, at 6 hrs, ∼90% of *ndd1-10A* cells already adapted but Rad53 was not dephosphorylated, in contrast to the normal expectation after adaptation (Lee et al., 2000, Pellicioli et al., 2001). This suggests that one important step in adaptation driven by Rad53 dephosphorylation might be to relieve the Ndd1-dependent repression of the *CLB2* cluster. Alternatively, the *ndd1-10A* mutant possibly allowed adaptation and the bypass of the DDC in a way that might differ from canonical adaptation.

We were also interested in the shape of the adaptation curve in the *ndd1-10A* mutant, which was much steeper than in the *NDD1* control strain when the first cells started to adapt. To better visualize this, we plotted the distribution of adaptation events over time and confirmed that the distribution in the *ndd1-10A* mutant was narrower and more skewed toward the right-hand side (Fig. 2C). These results are consistent with the idea that the Ndd1-dependent transcriptional response following DDC activation, including the downregulation of *CDC5*, strongly contributed to restrain adaptation and to its timing heterogeneity. However, another mechanism prevented adaptation before ∼2 hrs, as revealed in the *ndd1-10A* mutant.

Strikingly, the *ndd1-10A* allele completely rescued the adaptation defect of *cdc5-ad* whereas a second copy of wild-type *NDD1* did not (fraction of adapted cells at 24 hrs: mean ± SD = 77.8 ± 2.7% vs 3.1 ± 1.2%, respectively) (Fig. 2B). Adaptation in *cdc5-ad ndd1-10A* cells was slightly slower (*t*_*50*_ = 5.74 ± 0.39 hrs, mean ± SD) and less efficient than in *CDC5 ndd1-10A* cells, but still accelerated even compared to the *CDC5 NDD1* strain.

We conclude that phosphorylation of Ndd1 by Rad53 in response to telomere dysfunction slows down adaptation kinetics, likely by downregulating Cdc5, and contributes to the adaptation-defect of the *cdc5-ad* mutant. It is not required for the initial checkpoint arrest, but is important for its maintenance after 2 hrs.

### The transcriptional regulation of the *CLB2* cluster is essential to prevent adaptation in *cdc5-ad*

Since the kinase activity of Cdc5-ad is not impaired (Cheng et al., 1998, Rawal et al., 2016), stabilization of Cdc5-ad in the *ndd1-10A* mutant might be sufficient to allow adaptation. Alternatively, the checkpoint arrest might be bypassed in the *ndd1-10A* mutant by relieving the global transcriptional inhibition of the *CLB2* cluster of genes, regardless of the *cdc5-ad* allele. To distinguish between these two possibilities, we tested whether overexpression of Cdc5-ad alone would be sufficient to rescue *cdc5-ad*’s adaptation defect. Since *CDC5* is an essential gene, we examined adaptation in diploid strains in which expression of only one copy of Cdc5 or Cdc5-ad is under the control of a galactose-inducible (and glucose-repressible) promoter and tagged with 3xHA. Upon galactose addition, the 3xHA-tagged Cdc5 and Cdc5-ad were clearly overexpressed within an hour at both permissive and restrictive temperatures (Fig. 2D). We then performed microcolony assays to assess the adaptation ability of these strains, by spreading the cells on galactose-containing plates to induce Cdc5 or Cdc5-ad overexpression. We observed that overexpressing Cdc5 increased adaptation efficiency (Fig. 2E), likely by accelerating its kinetics as previously described (Vidanes et al., 2010). However, overexpressing Cdc5-ad was not sufficient to recover adaptation capacity (Fig. 2E). Thus, the adaptation defect of *cdc5-ad* was not simply due to a decreased expression and the Ndd1-dependent regulation of the *CLB2* cluster was important to prevent adaptation in the *cdc5-ad* mutant, beyond the repression of Cdc5-ad expression only.

### Cdc5-ad is hyperphosphorylated in a Mec1- and Tel1-independent manner

In addition to the transcriptional repression and degradation of Cdc5 and Cdc5-ad in response to telomere dysfunction, we observed that Cdc5-ad migrated as two forms in western blot (Fig. 1E), both of which were stabilized in *ndd1-10A* (Fig. 2A). To test whether the slow migrating band corresponds to a phosphorylated form of Cdc5-ad, we treated the samples with *λ*-phosphatase before electrophoresis and found that the slow migrating band was no longer present (Fig. 3A). Interestingly, at 3 hrs, both Cdc5 and the fast-migrating band of Cdc5-ad also displayed a slight shift in migration when treated with *λ*-phosphatase, suggesting that these forms were also phosphorylated. Since Cdc5 and Cdc5-ad seemed to be more phosphorylated after telomere dysfunction, we tested whether their phosphorylation depended on the DDC and particularly on Mec1 and Tel1. To do so, we induced telomere dysfunction in the hypomorph mutant *mec1-21* (Desany, Alcasabas et al., 1998, Sanchez, Desany et al., 1996) in the presence of nocodazole to prevent cycling beyond G2/M due to checkpoint deficiency. We found by western blot that in the *mec1-21* mutant, Rad53 was no longer hyperphosphorylated, as expected in this checkpoint-deficient mutant, but Cdc5-ad still migrated as two bands (Fig. 3B). Deletion of *TEL1* also did not affect the migration of Cdc5-ad as two bands after telomere dysfunction (Supplementary Fig. S2A). Thus, Cdc5-ad-specific phosphorylation did not depend on the Mec1 or the Tel1 branches of the DDC, when mutated separately. However, these two mutations had different effects on Cdc5 or Cdc5-ad levels upon telomere dysfunction. Consistent with our results regarding the transcriptional regulation of *CDC5* and *cdc5-ad* by Rad53 and Ndd1, *mec1-21* cells behaved like the *ndd1-10A* mutant with respect to Cdc5 and Cdc5-ad levels, while *TEL1* deletion did not affect them. This is presumably because the contribution of Tel1 to the DDC signaling is minor compared to Mec1. Thus, phosphorylation of Cdc5-ad in response to DNA damage was not dependent on Mec1 or Tel1.

**Figure 3.**
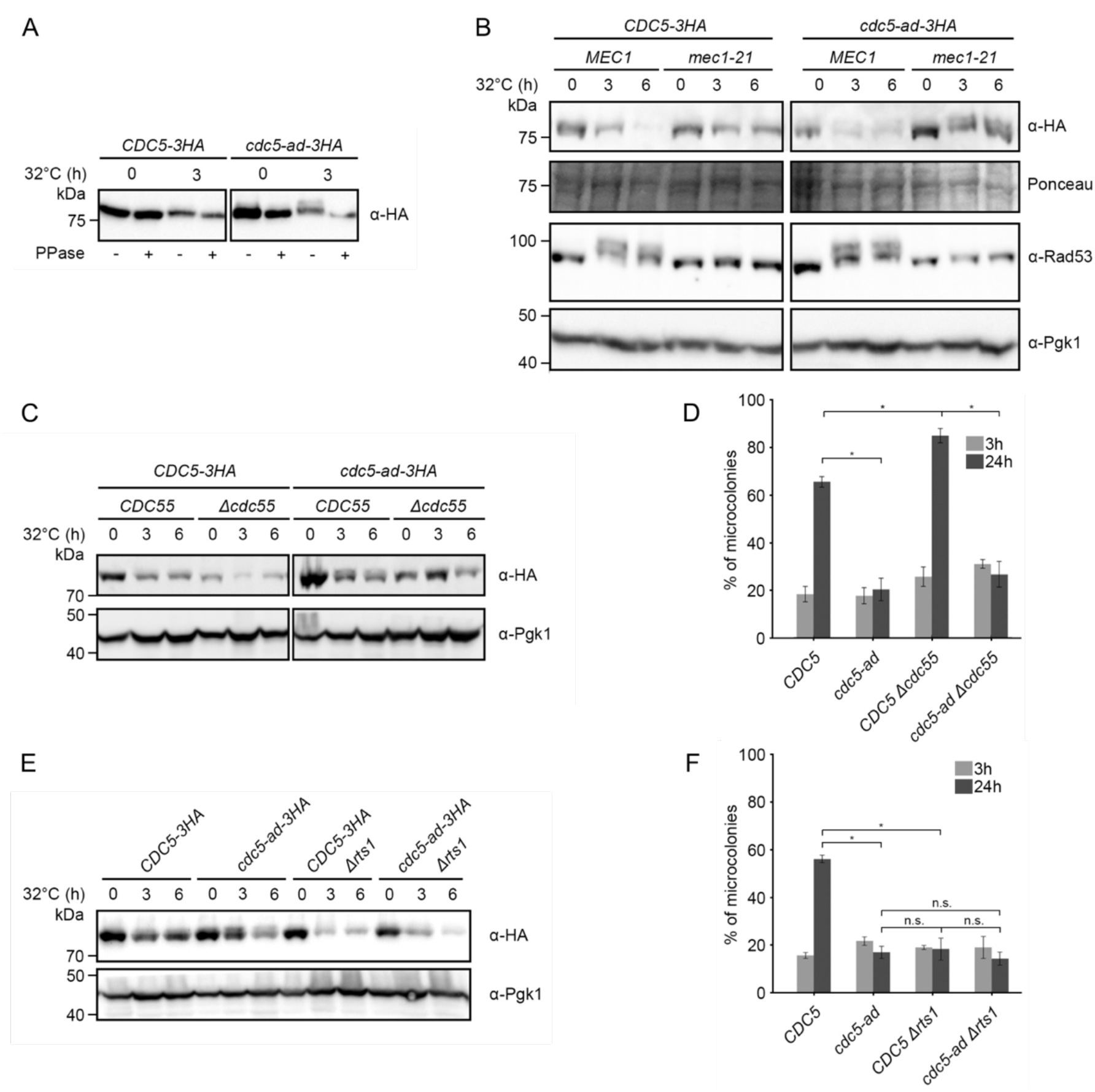
Cdc5-ad is phosphorylated in response to telomere dysfunction in a Mec1- and Tel1-independent and PP2A-dependent manner (see also Supplementary Fig. S2A). (**A**) Representative western blot of Cdc5 or Cdc5-ad from *cdc13-1* cells incubated at 32°C for the indicated amount of time. Samples were treated (+) or not (-) with *λ* phosphatase (PPase). (**B**) Representative western blot of Cdc5 or Cdc5-ad and Rad53 in response to telomere dysfunction in a wild-type *MEC1* or *mec1-21* mutant strain, incubated at 32°C for the indicated amounts of time, in the presence of nocodazole. (**C**) Representative western blot of Cdc5 or Cdc5-ad in a wild-type *CDC55* or *Δcdc55* mutant strain. (**D**) Microcolony assay measuring the fraction of microcolonies formed at 3 and 24 hrs in the indicated strains. Data are presented as means ± SD of N = 3 independent experiments. n ≥ 150 cells for each condition. (**E**) Representative western blot of Cdc5 or Cdc5-ad in a wild-type *RTS1* or *Δrts1* mutant strain. (**F**) Microcolony assay measuring the fraction of microcolonies formed at 3 and 24 hrs in the indicated strains. Data are presented as means ± SD of N = 3 independent experiments. n ≥ 150 cells for each condition.

### PP2A phosphatases affect adaptation in multiple ways

Since Cdc5 and Cdc5-ad were phosphorylated, particularly in response to telomere dysfunction and more so for Cdc5-ad, we wondered whether phosphatases regulate adaptation. In a recently published time-resolved phosphoproteomic analysis of mitosis (Touati, Hofbauer et al., 2019), residues T70, T238 and T242 in Cdc5 were found to be more phosphorylated in a PP2A^Rts1^ phosphatase mutant (*Δrts1*) in G2. But dephosphorylation of these residues after mitosis was not delayed, indicating that other phosphatases acted on them. In a PP2A^Cdc55^ phosphatase mutant (*Δcdc55*), dephosphorylation of residue S2 of Cdc5 was delayed in mitosis. To investigate the potential role of *RTS1* and *CDC55* in adaptation, we deleted these genes in our strains and assessed their adaptation phenotype.

Deletion of *CDC55* decreased the overall amount of Cdc5 and Cdc5-ad (Fig. 3C). Surprisingly, Cdc5-ad no longer migrated as two forms in *Δcdc55*, suggesting an indirect effect of PP2A^Cdc55^ on Cdc5-ad phosphorylation. In a microcolony assay, we found that the absence of *CDC55* slightly improved the adaptation level in *CDC5 cdc13-1* cells incubated at restrictive temperature (Fig. 3D), which was surprising given the decreased quantity of Cdc5 in this strain. This result suggested that Cdc55 counteracted adaptation and that adaptation might normally involve the inhibition or the bypass of Cdc55’s function(s). In the *cdc5-ad cdc13-1* strain, however, deletion of *CDC55* did not rescue the adaptation defect, indicating that the adaptation function affected in *cdc5-ad* is independent of Cdc55.

Surprisingly, *RTS1* deletion inhibited adaptation as strongly as the *cdc5-ad* mutant (Fig. 3F). The double mutant *Δrts1 cdc5-ad* was undistinguishable from either *Δrts1* and *cdc5-ad* with respect to adaptation. In the microcolony experiments, *Δrts1 cdc13-1* cells robustly arrested in G2/M after 3 hrs at 32°C, suggesting that the DDC was fully functional. Furthermore, *Δrts1* cells were proficient in recovery as they grew as well as *RTS1* cells in an assay where cells were subjected to a transient telomere dysfunction (for 10 hrs) before returning to a permissive temperature (Supplementary Fig. S2B). Therefore, *Δrts1* is a new bona fide adaptation mutant. We examined Cdc5 and Cdc5-ad proteins in the *Δrts1* mutant by western blot and found that both proteins started with a similar level but decreased in quantity faster than in a wild-type *RTS1* background (Fig. 3E). Interestingly, in the *Δrts1* mutant, Cdc5-ad did not migrate as two bands (Fig. 3E), indicating that Rts1 is important for Cdc5-ad’s phosphorylation. As for PP2A^Cdc55^, since PP2A^Rts1^ is a phosphatase, this effect must be indirect. We conclude that both phosphatases are involved in adaptation and that the hyperphosphorylation of Cdc5-ad is not essential to prevent adaptation.

### Phosphorylation of specific residues in Cdc5 modulates adaptation

To identify the phosphorylated residues in Cdc5 and Cdc5-ad, we immunoprecipitated Cdc5-3HA and Cdc5-ad-3HA in a *cdc13-1* mutant incubated at restrictive temperature for 0 or 3 hrs, using an anti-HA antibody, and performed mass spectrometry. Since the 3-hr samples had less Cdc5 and Cdc5-ad proteins (Fig. 1A and E), we took advantage of the *ndd1-10A* mutant, in which background both proteins were more abundant at this time point, to perform the experiment. The immunoprecipitated samples were separated by SDS polyacrylamide gel electrophoresis and silver stained (Supplementary Fig. S3A and B). The bands corresponding to Cdc5 and Cdc5-ad were cut from the gel. Proteins were extracted, trypsinized and analyzed by LC MS/MS. Fig. 4A and B summarize the phosphosites that we detected in two independent experiments.

**Figure 4.**
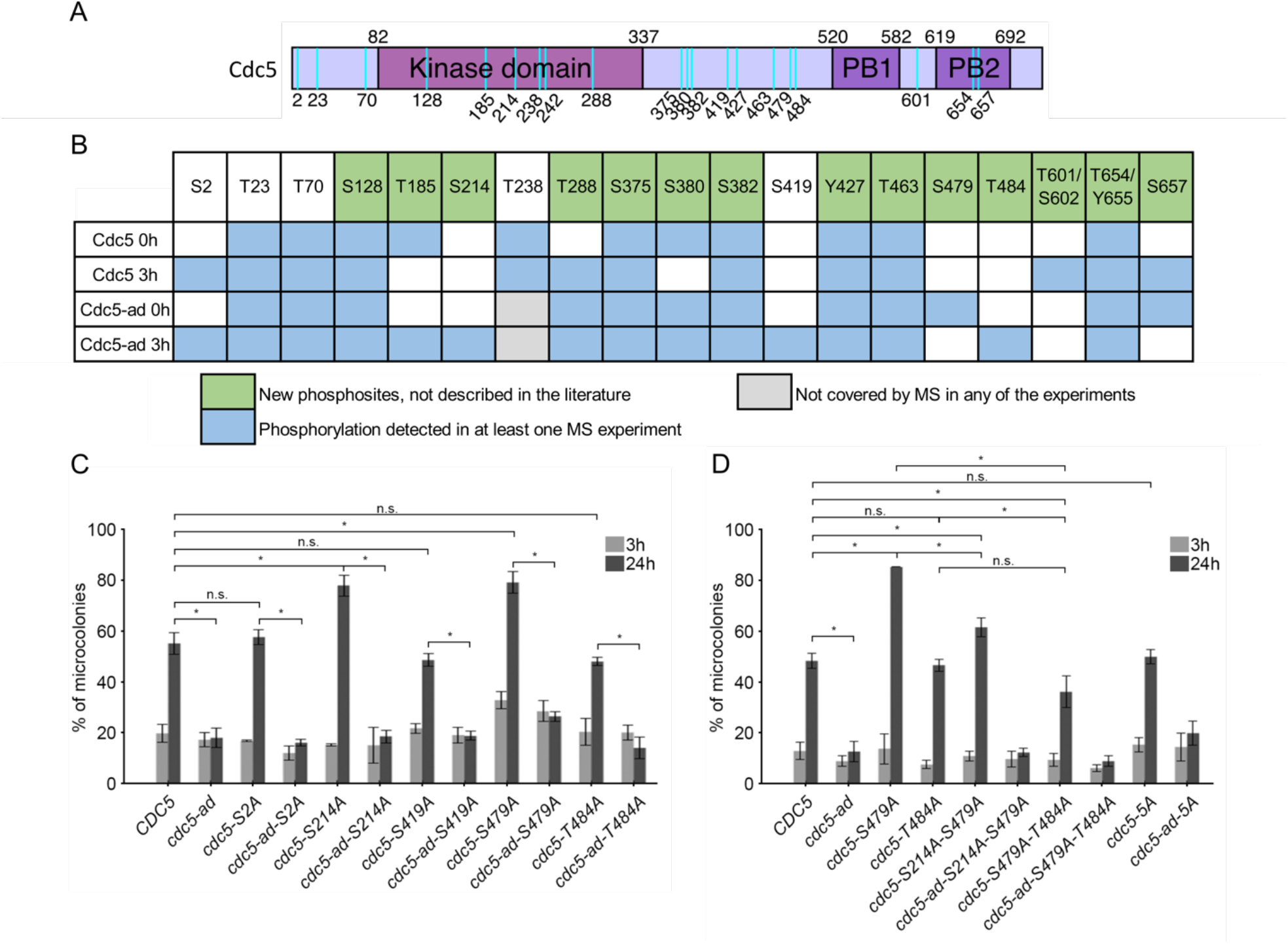
Phosphorylation of Cdc5 modulates adaptation. (**A**) Schematic representation of Cdc5 and its domains (kinase and Polo box domains PB1 and PB2) with the detected phosphorylation sites mapped (cyan vertical lines). (**B**) Summary table of phosphorylation sites identified by mass spectroscopy for each condition. For T601/S602 and T654/Y655, peptide analysis did not allow discrimination between the two putative phosphorylation sites. (**C**) Microcolony assay measuring the fraction of microcolonies formed at 3 and 24 hrs in strains carrying mutations to alanines at the residues S2, S214, S419, S479 and T484. Data are presented as means ± SD of N ≥ 3 independent experiments. n ≥ 150 cells for each condition. **(D)** Microcolony assay measuring the fraction of microcolonies formed at 3 and 24 hrs in strains carrying the indicated mutations, including double mutants S214A-S479A, S479A-T484A and all 5 sites mutated into alanines (“5A”). Data are presented as means ± SD of N ≥ 3 independent experiments. n ≥ 150 cells for each condition.

We found that Cdc5 and Cdc5-ad were heavily phosphorylated *in vivo*, in particular in DDC-arrested cells. Among the 19 phosphosites we detected in at least one experiment, 14 have not been described before in the literature, to our knowledge. The phosphosites mapped to all defined domains of Cdc5, including the N-terminus, the kinase domain and the Polo-box domain (PBD) (Fig. 4A).

We were particularly interested in residues that were differentially phosphorylated between Cdc5 and Cdc5-ad or between time points 0 and 3 hrs. We therefore focused on residues S2, S214, S419 (a minimal Cdk1 site), S479, and T484. We mutated each of these residues to alanine by Cas9-mediated mutagenesis and assessed the adaptation phenotype of the resulting mutant strains. The *cdc5-S214A* and *cdc5-S479A* mutants showed significantly increased adaptation level compared to wild-type, suggesting that phosphorylation of these sites inhibited adaptation (Fig. 4C). Other phosphosite mutants did not significantly alter adaptation level (Fig. 4C). Analysis of the double mutant *cdc5-S214A-S479A* showed that, while it still adapted better than the wild-type *CDC5*, it did not further increase adaptation and showed rather less efficient adaptation compared to each individual mutant (Fig. 4D). Because of the proximity of residues S479 and T484, we tested the effect of their combined mutation. Interestingly, the double mutant S479A T484A had an adaptation level comparable to the single T484A mutant, which was significantly lower than the high adaptation efficiency of S479A. Phosphorylation at S479 and T484 therefore appeared to have opposite effects, with phosphorylation at T484 being particularly important for efficient adaptation. While all 5 residues were phosphorylated in *cdc5-ad*, their combined mutation into alanines was not sufficient to alter the double-band migration profile in western blot (Supplementary Fig. S3C) and did not restore adaptation in the *cdc5-ad* mutant (Fig. 4D).

Overall, our results suggested that some of these residues, either by themselves or in combination, were involved in adaptation and that their phosphorylation modulates adaptation efficiency or timing. However, the adaptation deficiency of *cdc5-ad* did not solely depend on these phosphorylated residues.

### Cdc5’s functions in different pathways are important for adaptation

Since Cdc5 regulates multiple late cell cycle events, any of these could potentially be involved in Cdc5’s role in adaptation. For instance, the deletion of *BFA1*, the gene encoding the mitotic exit substrate of Cdc5, rescues the adaptation defect of *cdc5-ad* (Rawal et al., 2016). Cdc5’s interaction with other substrates, such as the RSC complex, is also important for adaptation, as evidenced by the adaptation defect of the *cdc5-16* mutant, which is mutated at 3 amino acids in the PBD domain, preventing its canonical interaction with primed substrates, including the RSC complex (Ratsima et al., 2016). We therefore asked whether these two *CDC5* mutants, *cdc5-ad* and *cdc5-16*, had the same underlying molecular defect in adaptation or whether they are affected in distinct pathways. We tested their genetic interaction by introducing these two alleles in diploid *cdc13-1/cdc13-1* cells and assessed the adaptation efficiency of this strain by microcolony assay (Fig. 5). As a control, we also combined the mutations of *cdc5-16* with the point mutation (L251W) of *cdc5-ad* to generate the *cdc5-ad-16* mutant.

**Figure 5.**
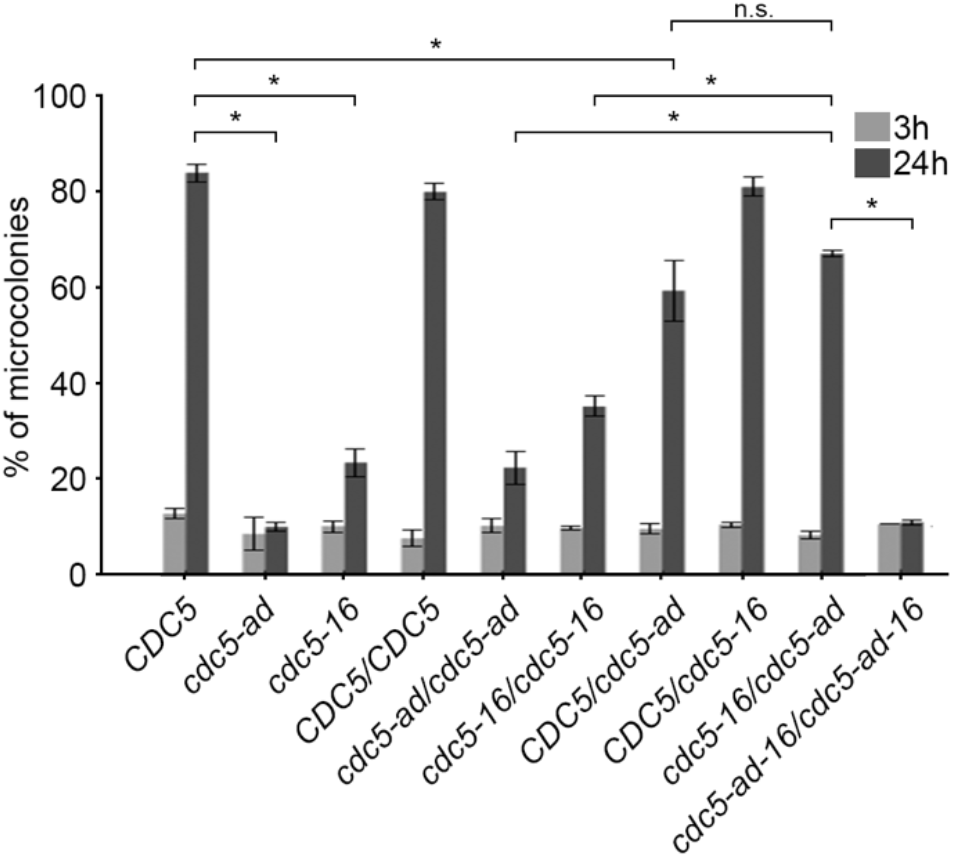
Mutant alleles *cdc5-16* and *cdc5-ad* complement each other in adaptation. Microcolony assay measuring the fraction of microcolonies formed at 3 and 24 hrs in the indicated mutants. Data are presented as means ± SD of N = 3 independent experiments. n ≥ 150 cells for each condition.

A wild-type copy of *CDC5* was sufficient to rescue the adaptation defect of *cdc5-ad* and *cdc5-16*. Strikingly, *cdc5-ad* and *cdc5-16* were able to complement each other’s adaptation deficiency as the heterozygous *cdc5-ad/cdc5-16* was adaptation proficient (Fig. 5). In contrast, the *cdc5-ad-16/cdc5-ad-16* mutant was as adaptation-deficient as *cdc5-ad/cdc5-ad*. We conclude that *cdc5-ad* and *cdc5-16* are affected in adaptation in different pathways or at least at different steps of the same pathway.

## Discussion

### Regulation of Cdc5 protein level in response to telomere dysfunction

Cdc5 was identified among the downstream targets of the DDC and proposed to be inhibited to prevent anaphase entry and mitotic exit in response to DNA damage (Sanchez et al., 1999). While a Rad53- and Ndd1-dependent inhibition of *CDC5* transcription was reported previously (Edenberg et al., 2014, Gasch et al., 2001, Jaehnig et al., 2013), how Cdc5 protein levels are regulated in response to telomere dysfunction has not been clearly established, especially as a function of time. Indeed, the temporal dimension of Cdc5 regulation is particularly important with respect to adaptation, which occurs late after the initial damage (4-16 hours) but depends on Cdc5 activity.

Here, we find that Cdc5 protein level decreases progressively in G2/M arrested cells after telomere dysfunction in a Mec1-, Rad53- and Ndd1-dependent manner, consistent with a transcriptional repression of *CDC5* expression. Incidentally, after telomere dysfunction, while Cdc5 degradation still relies on the proteasome, the previously described APC/C-Cdh1-dependent ubiquitinylation of Cdc5 KEN box and destruction box 1 (Arnold et al., 2015, Charles et al., 1998) is not involved. In a normal cell cycle, APC/C-Cdh1 starts to be active at the end of anaphase and in telophase and would not mediate Cdc5 degradation at the metaphase to anaphase transition. However, it was reported that Cdh1 is maintained in an active state in G2/M by the DDC in *cdc13-1* cells at restrictive temperature to prevent chromosome segregation (Zhang et al., 2009). The question of the mechanisms underlying Cdc5 degradation in this context, and whether Cdh1 is implicated, thus remains to be investigated.

Cdc5 being essential for adaptation, it might be surprising to observe that Cdc5 degradation after telomere dysfunction follows a kinetics that is inversely correlated to that of adaptation. One possibility would be that late cell cycle events are normally executed with an excess of Cdc5 proteins, consistently with the fact that a kinase mutant (*cdc5-77*) with less than 2% of wild-type activity is still viable (Ratsima, Ladouceur et al., 2011). This excess ensures a robust cell cycle progression but might be incompatible with checkpoint arrest. The decrease in Cdc5 quantity after telomere dysfunction would then be an essential step to enforce cell cycle control by the DDC. The low amount of Cdc5 might still be enough to trigger adaptation though, even after a long delay of 8-12 hrs. Cdc5’s activity in these conditions would then be finely controlled by other mechanisms such as posttranslational modifications. In this model, alterations of Cdc5 levels would tip the balance of Cdc5’s fine-tuned activity, thus explaining the many results correlating adaptation efficiency with the total amount of Cdc5, as we see in *ndd1-10A, Δrts1*, and in strains overexpressing Cdc5, or as described previously (Donnianni et al., 2010, Dotiwala et al., 2007, Hu et al., 2001, Vidanes et al., 2010). An exception to this rule would be the *Δcdc55* mutant in which adaptation is promoted while Cdc5 level is lower than in wild-type cells. In this case, the role of Cdc55 in preventing sister chromatid separation and inhibiting mitotic exit in the presence of DNA damage (Liang & Wang, 2007, Tang & Wang, 2006) might explain the slightly more efficient adaptation of the *Δcdc55* mutant despite lower Cdc5 level.

Alternatively, adaptation might be triggered by a transient burst of Cdc5 activity or quantity at the single cell level. Such a transient increase in activity or quantity would not be detected by measuring the total amount of Cdc5 at the population level, given the variability of adaptation timing at the single cell level. The transient and heterogeneous behaviour of Cdc5 expression would be consistent with the positive feedback regulation of Cdc5 on Ndd1, whereby Cdc5 phosphorylates Ndd1 on residue S85 to promote its binding to the promoter of the *CLB2* cluster of genes, thus stimulating its own expression (Darieva, Bulmer et al., 2006). However, while this possibility is worth exploring, the fact that most adaptation events are concentrated in the 8-12 hrs window (Fig. 2C) makes it unlikely that no global increase in Cdc5 quantity would be picked up by western blot.

A third possibility would be that Cdc5 functions early after the initial damage when its level is still high but its actual effect on adaptation takes several hours, potentially due to signal transduction through one or several pathways and to mechanisms that temporarily block adaptation progression (see below). There would then be a large delay between Cdc5’s peak activity after DNA damage and adaptation timing. This scenario would also be compatible with the observations correlating Cdc5 dosage with adaptation rate and efficiency.

### Post-translational modifications of Cdc5

Phosphorylation of Cdc5 and of Polo kinases in general is important for their kinase activity (Cheng et al., 1998, Hamanaka, Smith et al., 1995, Li, Wang et al., 2019, Mortensen et al., 2005, Qian, Yu et al., 1998, Rawal et al., 2016, Rodriguez-Rodriguez et al., 2016, Tavares, Glover et al., 1996). Additionally, Cdc5 is specifically phosphorylated in response to DNA damage in a Mec1- and Rad53-dependent manner (Cheng et al., 1998, Zhang et al., 2009). We also find that Cdc5 is phosphorylated after telomere dysfunction but most strikingly, we report that in this context Cdc5-ad migrates as two bands with distinct electrophoretic mobility. We note that, although it is induced by telomere deprotection, the slow migrating form of Cdc5-ad does not depend on Mec1 and Tel1 and is not directly due to the cells being in G2/M since a previous study showed that Cdc5-ad does not display an electrophoretic mobility shift during the cell cycle (Rawal et al., 2016).

Adaptation assays performed in mutants of individual phosphorylation sites or combinations of them to alanines indicate that they participate in the modulation of adaptation. For instance, phosphorylation of T484 likely promotes adaptation, whereas phosphorylation of S214 and S479 seems to have the opposite effect. In mutants of two very close residues, T484A appears to be dominant over S479A. We suggest that a complex combinatorial pattern of phosphorylations regulates Cdc5 function in adaptation, either by affecting its activity or its interaction with binding partners, including substrates. None of the mutations we tested or their combinations rescued, even partially, the adaptation defect of *cdc5-ad*. Even the mutation of 5 selected residues did not affect the adaptation phenotype of *cdc5-ad* nor the electrophoretic migration pattern of Cdc5-ad. We speculate that either phosphorylation of Cdc5-ad is not causal for the adaptation defect (as supported by the results in *Δcdc55* and *Δrts1* mutants) or that other phosphorylations that were not covered by the peptides detected by MS also participate in the adaptation defect of *cdc5-ad*.

Since the dynamics of dephosphorylation of Cdc5 residues were recently shown to be slightly altered in PP2A phosphatases mutants (Touati et al., 2019), we asked whether these phosphatases played a role in the electrophoretic mobility of Cdc5 and Cdc5-ad and in adaptation. Remarkably, both *Δcdc55* and *Δrts1* affect Cdc5/Cdc5-ad abundance and phosphorylation, and adaptation, but in very different ways. The mechanisms underlying the regulation of Cdc5/Cdc5-ad levels by the PP2A phosphatases are currently unknown. In *Δrts1*, the faster decrease of Cdc5 protein level results in a strong adaptation defect without affecting recovery after transient telomere dysfunction. In contrast, despite lower Cdc5 level, *Δcdc55* cells adapt slightly more efficiently, probably because it is necessary to bypass the functions of Cdc55 in inhibiting Cdc14 nucleolar release (Cdc-Fourteen Early Anaphase Release, FEAR) and in preventing chromosome segregation (Clift, Bizzari et al., 2009, Queralt, Lehane et al., 2006, Yaakov, Thorn et al., 2012). Importantly, since *Δcdc55 cdc5-ad* cells do not adapt, we infer that *cdc5-ad* is not defective in these processes after DNA damage or at least not only in these. Strikingly, the slow migrating form of Cdc5-ad is not present in the absence of *CDC55* or *RTS1*, suggesting an indirect regulation of Cdc5-ad phosphorylation by PP2A phosphatases, again showing the complexity of the post-translational regulation of Cdc5.

### Multiple Cdc5-dependent pathways control adaptation

Cdc5 is involved in many cell-cycle related processes (Botchkarev & Haber, 2018). Their coordination is important not only for a normal cell cycle but also during adaptation to ensure correct cell division despite persistent damage. Which of them is the limiting one or the first to be unblocked in adaptation and whether multiple Cdc5-dependent processes need to be overcome for adaptation remain unclear. Several observations made in our study, together with insights from literature, point to the idea that adaptation depends on the coordination of multiple pathways and events, often orchestrated by Cdc5.

In the first studies characterizing the DDC status during adaptation, Rad53 phosphorylation and kinase activity were shown to mirror cell cycle arrest and adaptation correlated well with the return of Rad53 to an unphosphorylated state (Lee et al., 2000, Pellicioli et al., 2001). Consistently, in adaptation mutants including *cdc5-ad*, Rad53 activation persisted for as long as 24 hours (Pellicioli et al., 2001). Since then, two adaptation defective mutants of *CDC5, cdc5-16* and *cdc5-T238A*, have illustrated an uncoupling between Rad53 status and adaptation (Ratsima et al., 2016, Rawal et al., 2016). In both cases (particularly obvious in *cdc5-16* cells), Rad53 returns to an unphosphorylated state while the cells stay arrested for much longer. Here, on the other hand, we find that in the *ndd1-10A* mutant, most cells have adapted within 4-6 hours of telomere deprotection while Rad53 is still hyperphosphorylated. Rad53 dephosphorylation is therefore neither necessary nor sufficient for adaptation, although in a wild-type context, it is likely the most straightforward way for the cell to alleviate the checkpoint arrest during adaptation. A direct phosphorylation of Rad53 by Cdc5 might contribute to checkpoint deactivation (Schleker, Shimada et al., 2010, Vidanes et al., 2010).

The initial checkpoint arrest does not require a functional Ndd1-dependent regulation of the *CLB2* cluster, as it is still present in the *ndd1-10A* mutant, and represents a first “locking” mechanism, *i*.*e*. that needs to be bypassed for adaptation to occur. The effect of the *ndd1-10A* mutant also demonstrates that, downstream of Rad53 activation, the transcriptional regulation of the *CLB2* cluster of genes is important for the long-term maintenance of the checkpoint arrest after 2-3 hours and therefore represents another locking mechanism for adaptation.

We also suggest that Cdc55 restrains adaptation through its functions in the FEAR network and in preventing chromosome segregation (Queralt et al., 2006, Yaakov et al., 2012), both of which antagonize Cdc5’s role in these processes. As noted before, the adaptation defect of *cdc5-ad* is independent of Cdc55 and must be caused by another locking mechanism, possibly the failure to activate MEN (Rawal et al., 2016). Additionally, we show that two adaptation mutants of *CDC5, cdc5-ad* and *cdc5-16*, can complement each other, demonstrating that multiple pathways controlled by Cdc5 are blocked simultaneously during checkpoint arrest and that Cdc5 needs to act on them all.

In conclusion, our work shows that multiple levels of regulation of Cdc5 in response to telomere dysfunction are critical for long-term checkpoint arrest, which requires blocking several processes involved in cell-cycle progression. Conversely, restarting all of them in a coordinated manner during adaptation is a complicated task for the cell. It is therefore remarkable that Cdc5 is involved in many of these processes, if not all, and this supports the idea that Cdc5 orchestrates, during adaptation, the multiple late cell cycle events leading to mitosis.

## Material and Methods

### Yeast strains

All strains are from the W303 background (*ura3-1 trp1-1 leu2-3,112 his3-11,15 can1-100*) corrected for *RAD5* and *ADE2* (Supplementary Table 1). They all contain the *cdc13-1* allele and were grown at the permissive temperature of 23°C in YPD (yeast extract, peptone, dextrose) media and at 32°C to induce telomere dysfunction. Overexpression and deletion strains were created using PCR-based methods as described in (Longtine, McKenzie et al., 1998). Point mutations were performed using Cas9-mediated gene targeting as described in (Anand, Beach et al., 2017).

### Microcolony assay

Microcolony assays were performed using the *cdc13-1* mutant to assess adaptation, as described in (Toczyski, 2006). Briefly, telomere dysfunction was induced in exponentially growing cells by incubation at 32°C for 3 hrs. 100 μL of a culture at OD_600 nm_ = 0.1 were then plated on a prewarmed plate. Plates were visualized on a dissection microscope (MSM System 400, Singer Instruments) immediately at 3 hrs and at 24 hrs. At each position, the number of cell bodies were counted and microcolonies were defined as comprising ≥ 3 cell bodies.

### SDS-PAGE and western blot analysis

Aliquots of 5 × 10^7^ cells were harvested by centrifugation. The pellets were lysed in 0.2 M NaOH on ice for 10 min and proteins were precipitated by the addition of 50 μL of 50% trichloroacetic acid. The samples were centrifuged at 16,100 g for 10 min at 4°C and the pellets were resuspended in 4× Laemmli buffer and heated for 5 min at 95°C. Samples were separated in a denaturing 7.5% 37.5:1 polyacrylamide gel, and proteins were transferred to a nitrocellulose membrane (Amersham Protran 0.45 NC, GE Healthcare).

The membranes were stained with Ponceau Red and immunoblotted with anti-Rad53 primary antibody (EL7.E1, Abcam), which recognizes both the unphosphorylated and phosphorylated forms of Rad53, anti-Cdc5 (11H12 and 4F10, Medimabs), anti-HA primary antibody (3F10, Roche), and anti-Pgk1 primary antibody (22C5D8, Abcam). Blots were then incubated with a horseradish peroxidase-coupled secondary antibody, and the signal was detected using ECL reagent (Amersham, GE Healthcare). All western blots were independently replicated in the laboratory.

### Immunoprecipitation

Immunoprecipitation was performed as in (Cohen, Amiott et al., 2011) with minor modifications. After growth in the different conditions described in the results, ∼10^9^ cells were harvested, washed in Milli-Q water and resuspended in lysis buffer (50 mM Tris-HCl pH 8.0, 150 mM NaCl, 0.6% Triton X-100, 10% glycerol, protease inhibitors (cOmplete Mini EDTA-free Protease Inhibitor Cocktail, Roche) and phosphatase inhibitors (Phosphatase Inhibitor Cocktail Set II, Millipore). Cells were lysed with glass beads in a cooled FastPrep (MP Biomedicals). Cell lysates were cleared by centrifugation at 13 000 g for 30 min and supernatants were incubated for 30 min with 2.5 μg/sample anti-HA antibodies (clone 3F10, Roche) bound to 50 μL/sample (or 1.5 mg/sample) protein G-coated magnetic beads (Dynabeads Protein G, Invitrogen). The subsequent steps of immunoprecipitation and washes were performed according to the manufacturer’s instructions. The immunoprecipates were eluted in 2x Laemmli buffer.

### Mass spectrometry analysis

After separation by SDS-PAGE, proteins were fixed and revealed by silver nitrate staining according to the protocol described in (Rabilloud, 2012). Bands of interest were excised and prepared for trypsin digestion. Briefly, proteins were destained with a freshly prepared solution containing 15 mM potassium ferricyanide and 50 mM sodium thiosulfate. Proteins were reduced by 10 mM dithiotreitol in 50 mM ammonium bicarbonate (AMBIC) for 30 min at 56°C and further alkylated by incubation in the dark with 50 mM iodoacetamide in 50 mM AMBIC. Then, protein samples were digested overnight at 37°C with 125 ng modified porcine trypsin (Trypsin Gold, Promega). Peptides were extracted under acidic conditions and dried out using a speedvac concentrator. Then, peptide mixtures were resuspended in 10 μL of solvant A (0.1% (v/v) formic acid in 3% (v/v) acetonitrile) and frozen at -20 °C until use.

Mass spectrometry analyses were performed on a Q-Exactive Plus hybrid quadripole-orbitrap mass spectrometer (Thermo Fisher, San José, CA, USA) coupled to an Easy 1000 reverse phase nano-flow LC system (Proxeon) using the Easy nano-electrospray ion source (Thermo Fisher). Five microliters of peptide mixture were loaded onto an Acclaim PepMap™ precolumn (75 μm x 2 cm, 3 μm, 100 Å; Thermo Scientific) equilibrated in solvent A and separated at a constant flow rate of 250 nl/min on a PepMap™ RSLC C18 Easy-Spray column (75 μm x 50 cm, 2 μm, 100 Å; Thermo Scientific) with a 90 min gradient (0 to 20% B solvent (0.1% (v/v) formic acid in acetonitrile) in 70 min and 20 to 37% B solvent in 20 min).

Data acquisition was performed in positive and data-dependent modes. Full scan MS spectra (mass range m/z 400-1800) were acquired in profile mode with a resolution of 70,000 (at m/z 200) and MS/MS spectra were acquired in centroid mode at a resolution of 17,500 (at m/z 200). All other parameters were kept as described in (Perez-Perez, Mauries et al., 2017).

Protein identification was performed with the MASCOT software (version 4; Matrix Science, London, UK) via the Proteome discoverer software (version 2.2; Thermo Scientific) with *S. cerevisiae* UniprotKB protein database. The search parameters were set as followed: two missed cleavages allowed cysteine carbamidomethylation as fixed modifications and methionine oxidation, N Ter protein acetylation and phosphorylation on serine, tyrosine and threonine as variable modifications with a peptide mass tolerance of 10 ppm. FDR (False discovery rate) for protein identification was fixed at 0.01.

### Time-lapse microscopy

Exponentially growing cells at OD_600 nm_ = 0.2 were injected into the microfluidics chambers of a CellASIC plate (Y04C-02 plate for haploid cells, CellASIC ONIX2 Microfluidic System, Millipore), through the inlet wells. A constant flow of rich media (20 μL/hr for 0.036 μL chambers) fed the chambers, which were kept at the constant temperature of 32°C. Cells in the microfluidic device were imaged using a fully motorized Axio Observer Z1 inverted microscope (Zeiss) with a 100× immersion objective, a Hamamatsu Orca R2 camera, and constant focus maintained with focus stabilization hardware (Definite focus, Zeiss). The temperature was maintained at 32°C with a controlled heating unit and an incubation chamber. Images were acquired every 10 min using ZEN software (Zeiss). All aspects of image acquisition were fully automated and controlled, including temperature, focus, stage position, and time-lapse imaging.

## Acknowledgements/Funding

We are grateful to Jiri Veis, Gustav Ammerer and Jim Haber for providing plasmids and yeast strains. We thank Sandra Touati for helpful discussions and for sharing unpublished data, and Leticia Berkate for her technical help.

Research in DD’s laboratory is funded by a Foundation grant from CIHR (FDN–167265). DD and the University of Ottawa are also funded by the Canada Research Chair in Chromatin Dynamics and Genome Architecture. Work in MTT’s laboratory is supported by the “Fondation de la Recherche Medicale” (“Équipe FRM EQU202003010428”), by the French National Research Agency (ANR) as part of the “Investissements d’Avenir” Program (LabEx Dynamo ANR-11-LABX-0011-01) and ANR-16-CE12-0026, and The French National Cancer Institute (INCa_15192). H.C. was supported by a doctoral grant from the Paris Sciences et Lettres (PSL) Idex program implemented by the ANR (ANR-10-IDEX-0001-02 PSL). Research in ZX’s laboratory is supported by the ANR grant “AlgaTelo” (ANR-17-CE20-0002-01), by Ville de Paris (Programme Émergence(s)) and by the Émergence grant of Sorbonne Université. The Proteomic Platform of the IBPC (PPI) was supported by LABEX DYNAMO (ANR-LABX-011) and EQUIPEX (CACSICE ANR-11-EQPX-0008).

## Author contributions

Conceptualization: DD, MTT, ZX. Formal Analysis: HC, OI, JLP, MH, ZX. Investigation: HC, OI, JLP, MH. Supervision: MTT, ZX. Resources: DD, MTT, ZX. Writing – original draft: ZX. Writing – review & editing: all authors.

## Supplementary data

**Supplementary Table 1.**
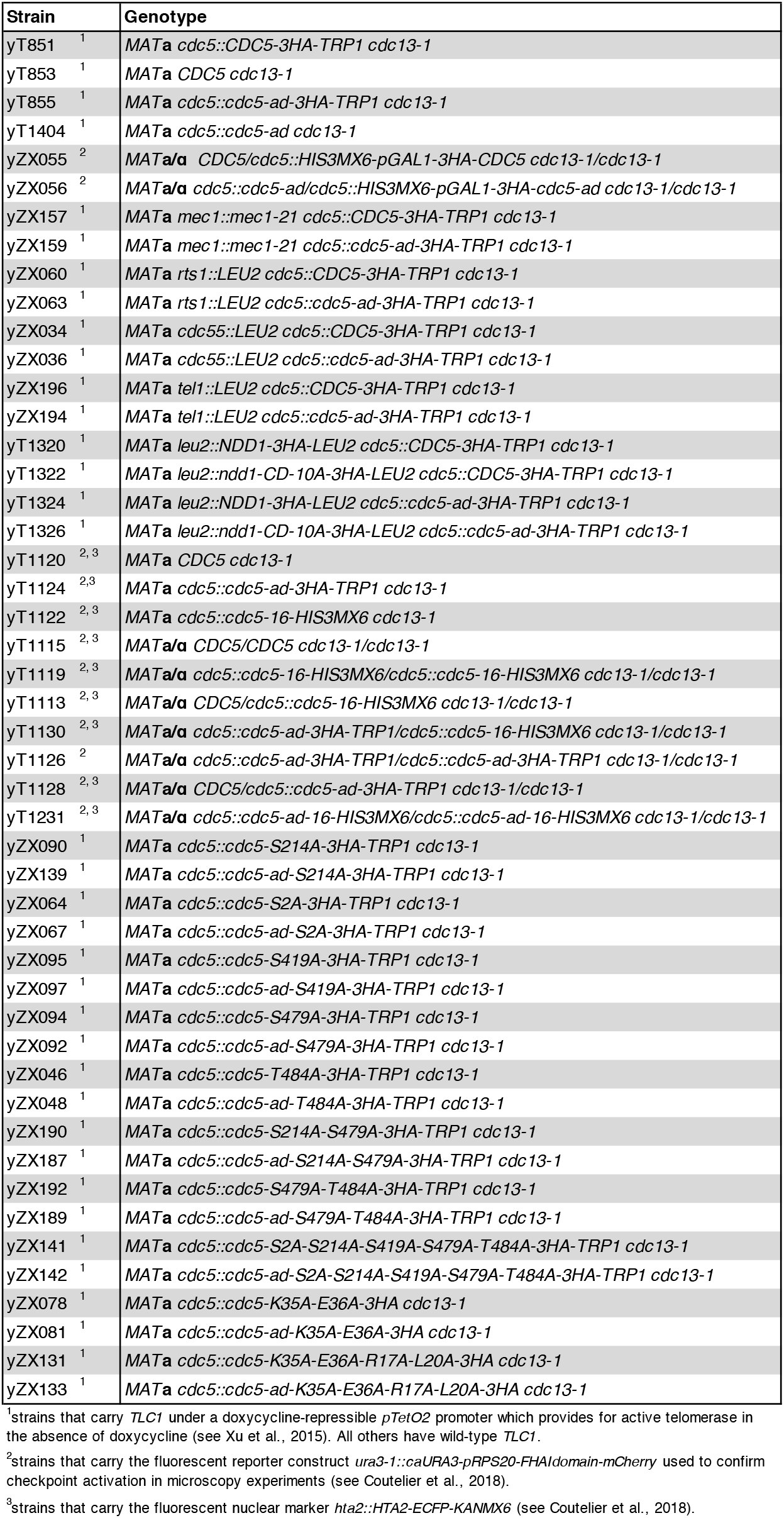
Yeast strains used in this study.

**Supplementary Figure S1.**
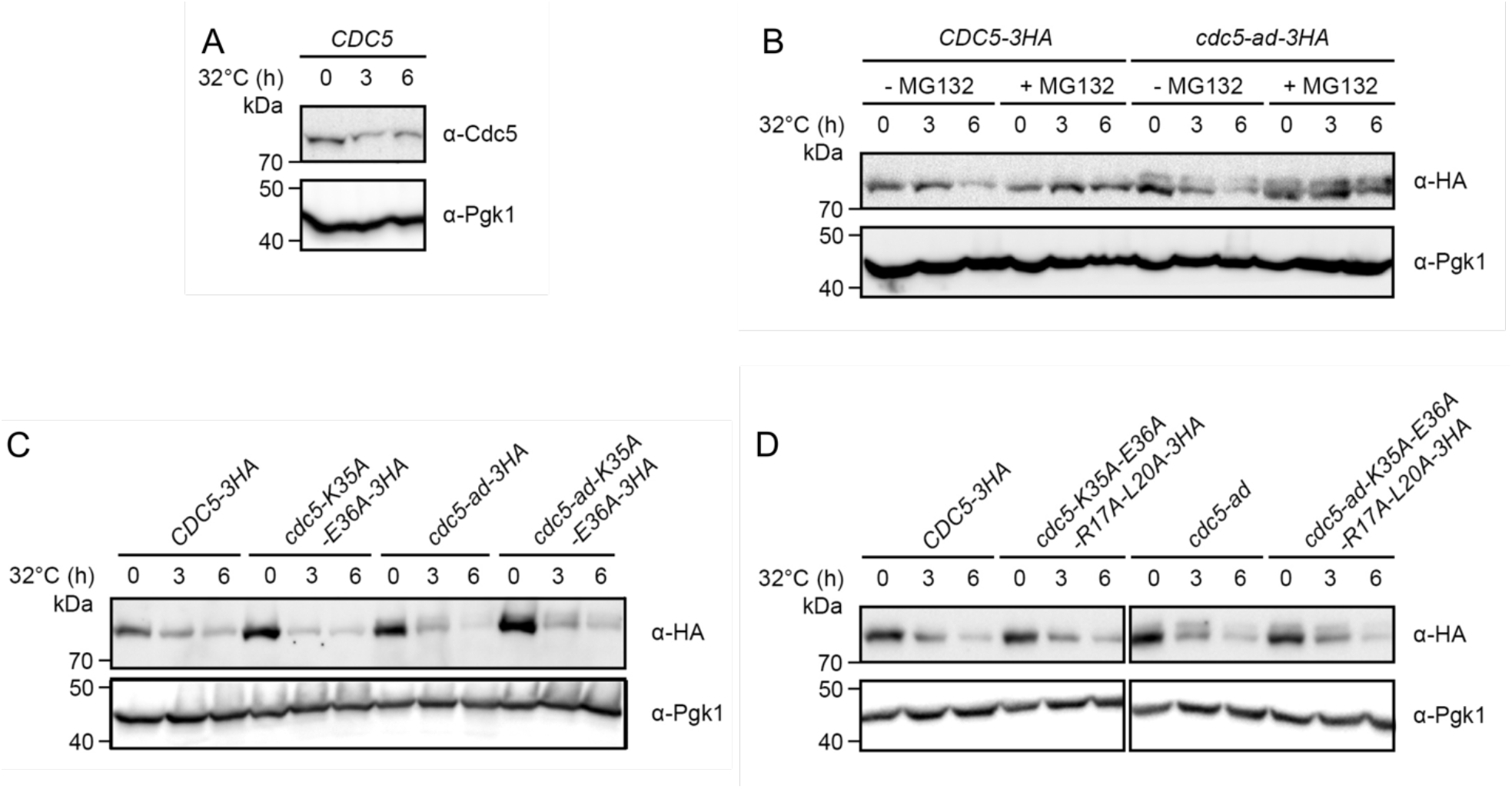
**(A)** Representative western blot of untagged Cdc5 in the *cdc13-1* mutant incubated for the indicated times at 32°C. Pgk1 is shown as a loading control. **(B)** Representative western blot of Cdc5 and Cdc5-ad in the *cdc13-1* mutant incubated for the indicated times at 32°C, with or without MG132. **(C)** Representative western blot of Cdc5 and Cdc5-ad, with or without additional K35A E36A mutations in the KEN box. The strains all contain the *cdc13-1* allele and were incubated for the indicated times at 32°C. **(D)** Representative western blot of Cdc5 and Cdc5-ad, with or without additional K35A E36A mutations in the KEN box and R17A L20A in the destruction box 1. The strains all contain the *cdc13-1* allele and were incubated for the indicated times at 32°C.

**Supplementary Figure S2.**
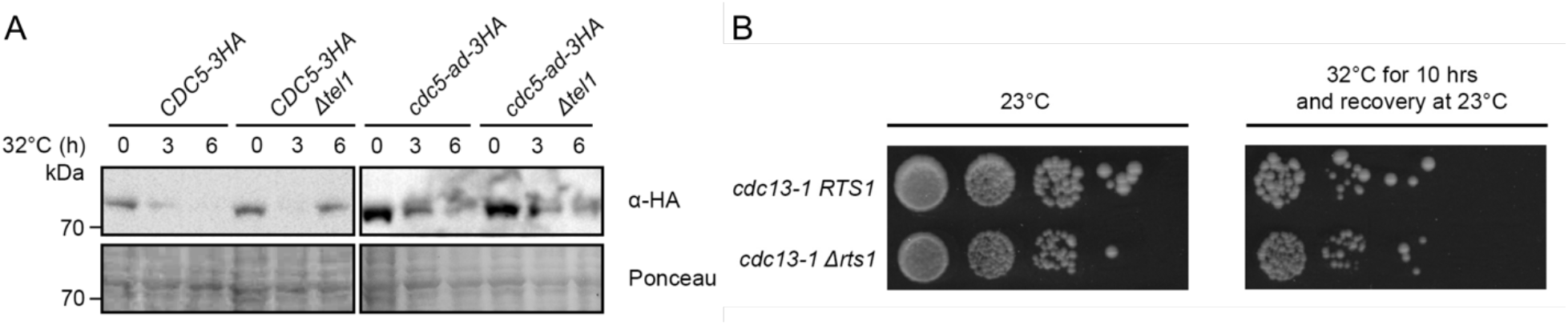
**(A)** Representative western blot of Cdc5 and Cdc5-ad in the *cdc13-1 TEL1* or *Δtel1* strains, incubated for the indicated times at 32°C, in the presence of nocodazole. The increase in Cdc5 protein level at 6 hrs compared to 3hrs in the *Δtel1* strain is reproducible. (B) Recovery assay in *cdc13-1 RTS1* or *Δrts1* strains. Ten-fold serial dilutions of cells were incubated at 32°C for 10 hrs to induce transient telomere dysfunction and allowed to recover at 23°C afterwards, or continuously grown at 23°C as a control.

**Supplementary Figure S3.**
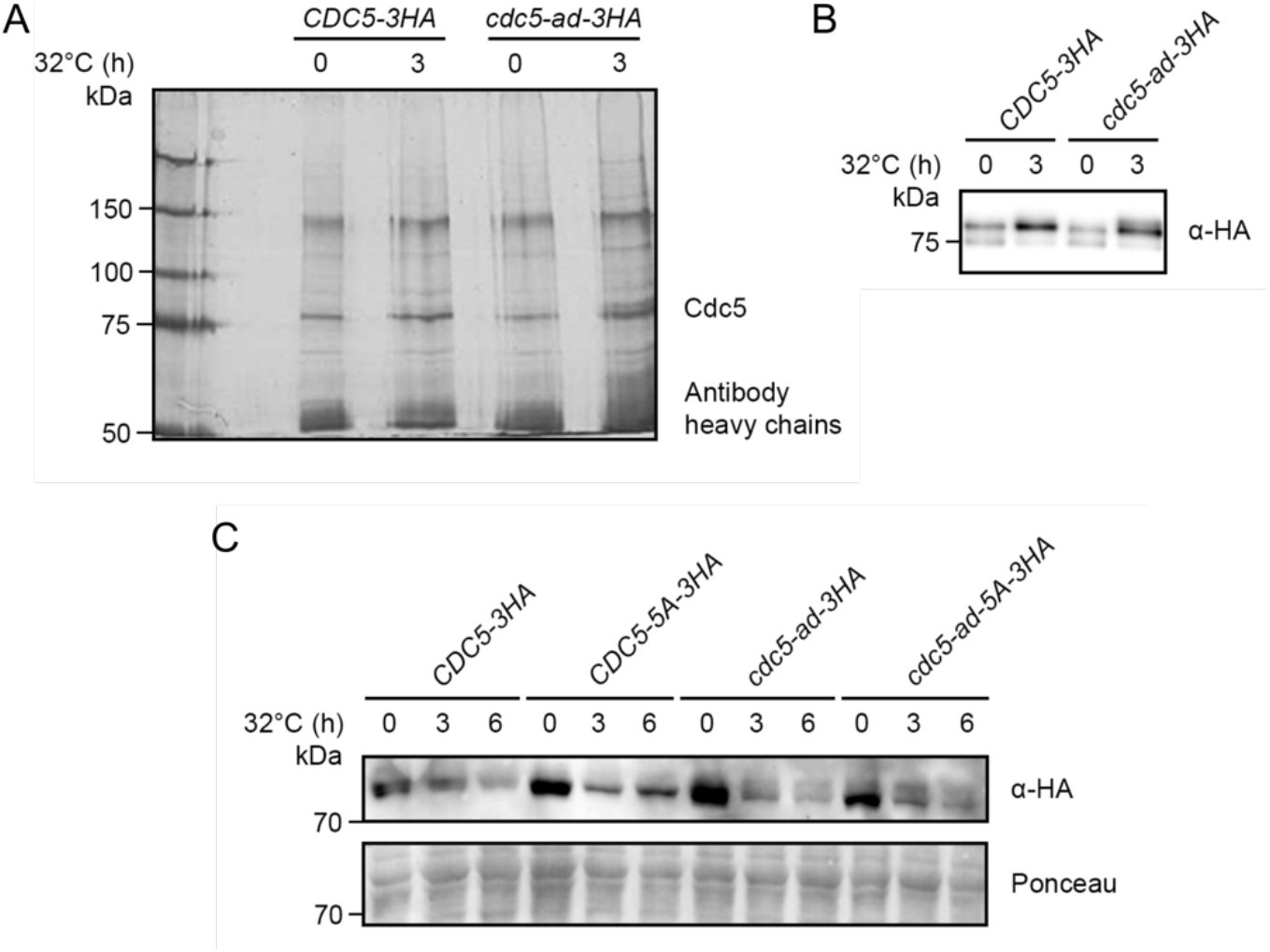
(**A**) Silver-nitrate-stained gel showing immunoprecipitation products from *ndd1-10A cdc13-1* strains incubated for the indicated amounts of time at 32°C, which were then cut out for mass spectroscopy analysis. **(B)** Western blot of the immunoprecipitated Cdc5 and Cdc5-ad from *ndd1-10A cdc13-1* strains incubated at 32°C for the indicated amount of time. **(C)** Representative western blot of Cdc5 and Cdc5-ad, with or without additional mutation of all 5 phosphosites into alanines (“5A”), in the *cdc13-1* strain, incubated for the indicated times at 32°C

